# Comparative Genomic Analysis of Fifty-Two *Staphylococcus aureus* Isolates Identified from Uncharacterized Staphylococcus Genomes in the NCBI database

**DOI:** 10.1101/2020.06.18.159814

**Authors:** Mohamed A. Abouelkhair

## Abstract

**Background:** *Staphylococcus aureus* is a major bacterial pathogen that causes a variety of diseases, ranging from wound infections to severe bacteremia or food poisoning. The course and severity of the disease are mainly dependent on the bacterium genotype as well as host factors. Whole-genome sequencing (WGS) is currently the most extensive genotyping method available, followed by bioinformatic sequence analysis.

**Methods:** A total of 253 uncharacterized staphylococcus genome sequences were downloaded from the National Center for Biotechnology Information (NCBI) (August 2012 to March 2020) from different studies. Samples were clustered based on core and accessory pairwise distances between isolates and then analyzed by multilocus sequence typing tool (MLST). Staphylococcal Cassette Chromosome *mec* (SCC*mec*), spa typing, variant calling, core genome alignment, and recombination sites prediction were performed on detected *S. aureus* isolates. *S. aureus* isolates were also analyzed for the presence of genes coding for virulence factors and antibiotic resistance.

**Results and conclusion:** Uncategorized genome sequences were clustered into 24 groups. About 182 uncharacterized Staphylococcus genomes were identified at the species level based on MLST, including 32 *S. lugdunensis* genome sequence, thus doubling the number of the publicly accessible *S. lugdunensis* genome sequence in Genbank. MLST identified another four species (*S. epidermidis* (33/253), *S. lugdunensis* (32/253), *S. haemolyticus* (41/253), *S. hominis* (24/253) *and S. aureus* (52/253)). Among the 52 *S. aureus* isolates, 21 (40.38%) isolates carried *mec*A gene, with 57.14% classified as SCC*mec* IV. The results of this study provide knowledge that facilitates evolutionary studies of staphylococcal species and other bacteria at the genome level.

## Introduction

*Staphylococcus aureus* bacterium is worldwide distributed and is responsible for several human and animal diseases ranging from mild to life-threatening infections (1, 2). It is of considerable importance because of its ability to induce a multitude of infections and adapt to various environmental conditions. *S. aureus* is one of the most significant causes of hospital and community-acquired infections, with serious consequences (3). The emergence of methicillin-resistant *S. aureus* (MRSA) places a major burden on the public health care system. Along with evolving several mechanisms that confer resistance to antibiotic compounds, *S. aureus* produces a vast arsenal of virulence proteins, including exotoxins, tissue-degrading enzymes, leucocidins, and immunomodulating proteins (4–6).

Accurate molecular typing is important for tracking outbreaks, determining the probable source of colonization (livestock or human associated), and distinguishing between the community and hospital-acquired strains. Various typing methods such as multilocus sequence typing (MLST), SCC*mec* typing, and spa typing can be used to identify methicillin-resistant *S. aureus* (MRSA) lineages and strains. Molecular biology and biotechnology advances have made the entire genome sequencing accessible for research in microbiology. Whole-genomic microbes’ sequences can provide comprehensive information on virulence factors, pathogenesis, drug resistance, metabolism, the interaction between host-pathogen, MLST, SCC*mec*, and spa types.

GenBank ® is an extensive and reliable database which contains publicly available nucleotide sequences (7). There are more than 14,000 genome assemblies available for 54 staphylococcus species. In this study, we used a genomic approach to identify *S. epidermidis*, *S. lugdunensis*, *S. haemolyticus*, *S. hominis*, *and S. aureus* from uncategorized genome sequences in the NCBI database. They are annotated as isolates in a staphylococcus population, which are significantly different from currently recognized species. The specific objectives of this study were to characterize these isolates and identify the genomic characteristics of identified *S. aureus* isolates.

## Material and Methods

### Genomes annotation, and population structure analysis

Uncharacterized staphylococcus genome sequences were downloaded from NCBI (https://www.ncbi.nlm.nih.gov/genome/browse#!/prokaryotes/13533/) (August 2012 to March 2020) from different studies. Enterobacter cloacae with a genome size of 4.8 Megabases (Mb) in addition to four plasmids were found in the group and excluded from the study.

The assemblies were annotated using Prokka (https://github.com/tseemann/prokka) (8). We used PopPUNK (Population Partitioning Using Nucleotide K-mers; https://poppunk.readthedocs.io/en/latest/) to elucidate the population structure of staphylococcus based on the divergence of both shared sequence and gene content within a population. PopPUNK compares all possible pairs of genomes by calculating the proportion of k-mers shared of different lengths to determine the distances between the core and accessory genomes. It then generates a scatterplot of the two distances to reveal the isolates predicted to cluster (9). The genome sequences of identified *S. aureus* were uploaded to the Type (Strain) Genome Server (TYGS), a free bioinformatics platform available under https://tygs.dsmz.de, for a whole genome-based taxonomic analysis (10). Determination of closest type strain genomes was done in two complementary ways: First, all *S. aureus* genomes were compared against all type strain genomes available in the TYGS database via the MASH algorithm, a fast approximation of intergenomic relatedness (11), and, the ten type strains with the smallest MASH distances chosen. Second, an additional set of ten closely related type strains was determined via the 16S rDNA gene sequences. These sequences were extracted from the *S. aureus* genomes using RNAmmer (12), and each sequence was subsequently BLASTed (13) against the 16S rDNA gene sequence of each of the currently 11820 type strains available in the TYGS database. This was used as a proxy to find the best fifty matching type strains (according to the bitscore) for each *S. aureus* genome and to subsequently calculate precise distances using the Genome BLAST Distance Phylogeny approach (GBDP) under the algorithm ‘coverage’ and distance formula *d_5_* (14). These distances were finally used to determine the ten closest type strain genomes for each of the *S. aureus* genomes.

All pairwise comparisons among the set of genomes were conducted using GBDP and accurate intergenomic distances inferred under the algorithm ‘trimming’ and distance formula *d_5_* (14). One hundred distance replicates were calculated each. Digital DDH values and confidence intervals were calculated using the recommended settings of the GGDC v.2.1 (14). The resulting intergenomic distances were used to infer a balanced minimum evolution tree with branch support via FASTME v.2.1.4, including SPR postprocessing (15). Branch support was inferred from 100 pseudo-bootstrap replicates each. The trees were rooted at the midpoint (16) and visualized with PhyD3 (17). The type-based species clustering using a 70% dDDH radius around each of the 28 type strains was done as previously described (10).

### *In silico* molecular typing

Sequence type (ST) identification of isolates was performed using the program MLST (https://github.com/tseemann/mlst) (18), which extracts the sequences of seven housekeeping genes (*arcC, aroE, glpF, gmk, pta, tpi, yqiL*) from the assembly files. This program made use of the PubMLST website (https://pubmlst.org/) developed by Keith Jolley (19) and sited at the University of Oxford.

### Detection of genes coding for antimicrobial resistance and virulence proteins

We screened all of the genomes for known resistance genes using NCBI Antimicrobial Resistance Gene Finder v.3.8 (AMRFinderPlus) (https://github.com/ncbi/amr) (20). To identify the gene coding for virulence factors, we used the contig-based search method ABRicate v.0.8.13 (https://github.com/tseemann/abricate). We created a custom abricate database for *staphylocoagulase*.

### SCC*mec* typing, spa typing

Genomes carrying the *mec*A-carrying chromosomal cassette SCC*mec* were identified using SCCmecFinder v.1.2 (21) with minimum thresholds of >60% for sequence coverage and >90% sequence identity whereas spa typing was performed using spaTyper v.1.0 (22). We used the default parameters for each program.

### Predicting Recombination Sites by Phylogenetic Analysis

We aligned the 52 isolates against the reference genome of *S. aureus* NCTC 8325 by using Snippy v.4.6.0 (https://github.com/tseemann/snippy), which uses the Burrow-Wheeler Aligner (BWA) to map the reads to the reference and then calls the subsequent single-nucleotide polymorphisms (SNPs) and insertions/deletions (indels). We used the whole-genome core SNPs alignment output from Snippy for downstream phylogenetic analysis, assessed recombination sites by using Genealogies Unbiased By recomBinations In Nucleotide Sequences v.2.4.1 (Gubbins) (https://github.com/sanger-pathogens/gubbins) (23). After five iterations, Gubbins reached a stable tree topology and determined regions of genetic recombination. Fastree v.2.1.10 were used to build a high-resolution phylogeny from the core SNP genome. We visualized the resulting phylogenetic tree, core genome SNPs, and recombination sites by using Phandango version v.1.3.0 (24).

### Code availability

All tools used for the analysis are publicly available and fully described in the “Method” sections.

## Results

### Multiple Staphylococcus species identified among unnamed staphylococcus isolates

A total of 253 uncharacterized staphylococcus genome sequences were downloaded from NCBI from August 2012 to March 2020 (median total length: 2.53498 megabases, median protein count: 2361 and median GC%: 32.7).

Twenty-four clusters were identified based on core and accessory pairwise distances between isolates using MinHash-optimized k-mer comparisons **Figure. 1**.

**Figure.1:**
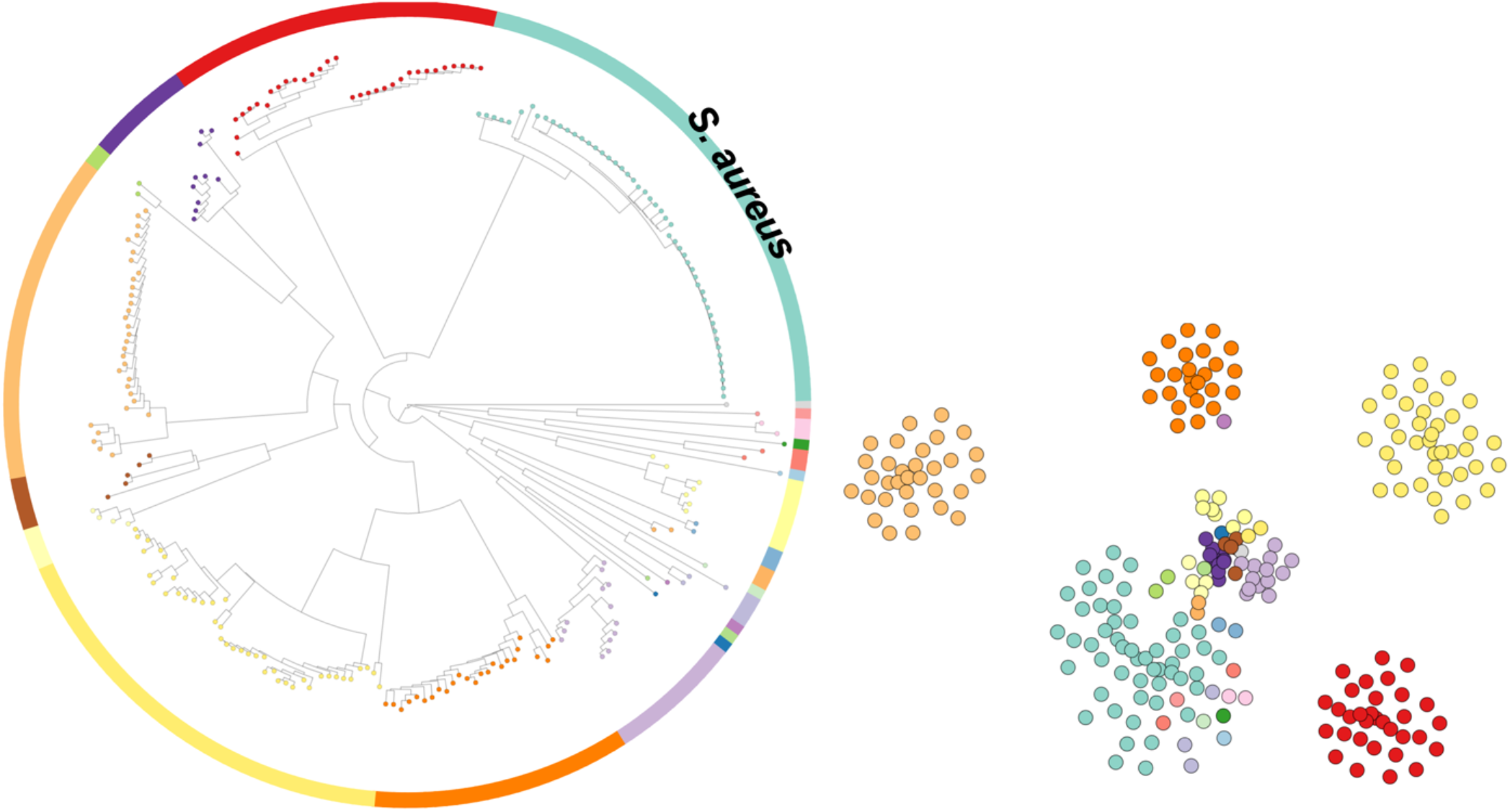
Twenty-four clusters were identified based on core and accessory pairwise distances between isolates using MinHash-optimized k-mer comparisons. Population clusters were visualized using microreact.

MLST identified five species (*S. epidermidis* (33/253), *S. lugdunensis* (32/253), *S. haemolyticus* (41/253), *S. hominis* (24/253) *and S. aureus* (52/253)). Fifty-six isolates were not typed by MLST (20.94%). Forty-five *S. aureus* genomes categorized into seven clonal complexes (CC1, CC5, CC8, CC15, CC30, CC45, and CC97). Among those, twenty-two genomes belong to clonal complex CC5 (ST5; n=21 genomes and ST105; n=1 genome) and fourteen genomes grouped in CC8 (ST8; n=13 genomes and ST72; n=1 genome) **Figure.2**. MLST typing and clonal complex of *S. aureus* are listed in **Table S1**. MLST typing and allele profiles of the other four species are listed in **Table S2**.

**Figure.2:**
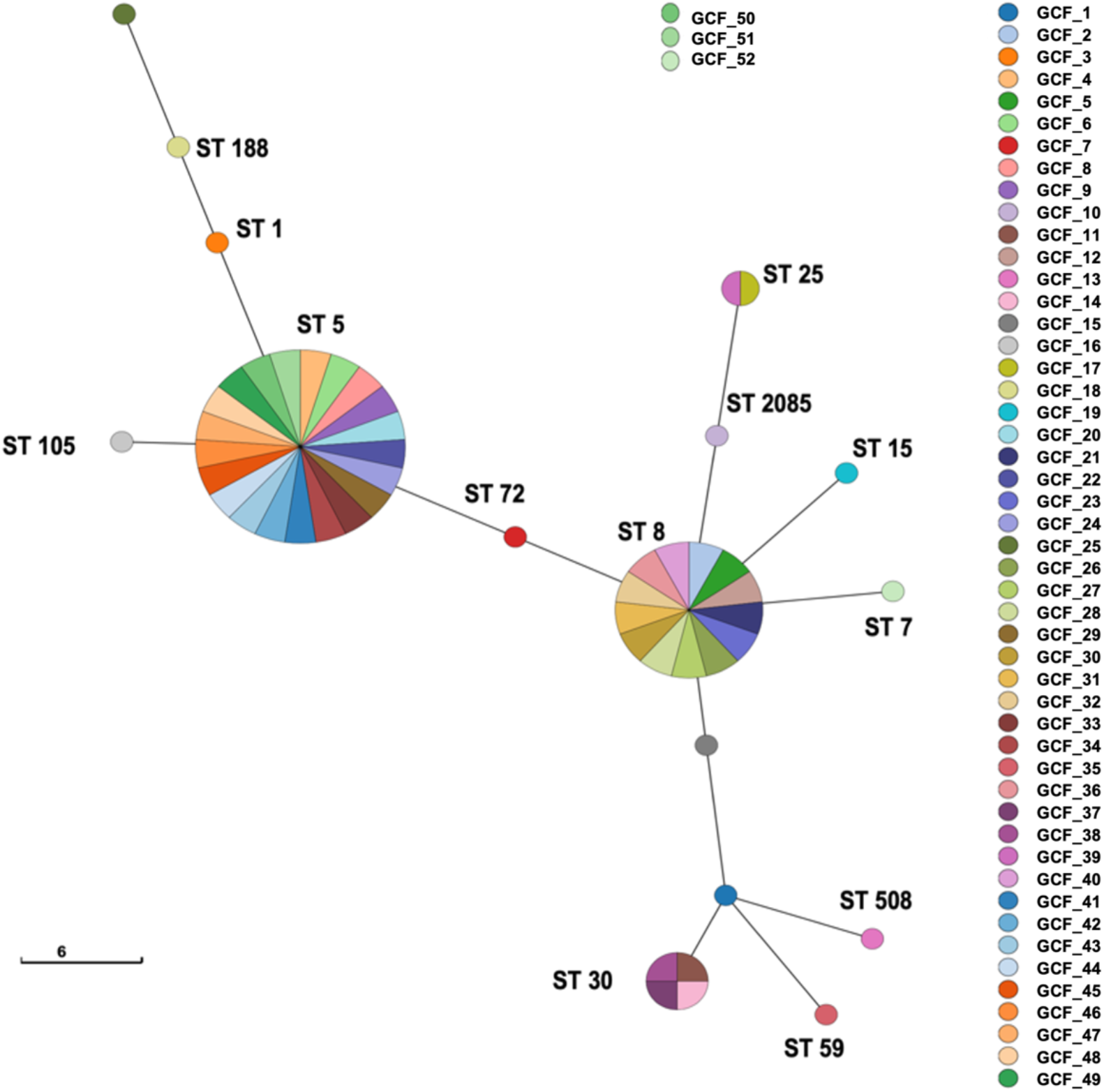
MLST typing of *S. aureus* population. The data visualized using GrapeTree (25) and nodes were colored based on sample ID.

Type-based species clustering yielded 22 species clusters, within the staphylococcus genus, and the *S. aureus* isolates were assigned to the same cluster of *S. aureus* DSM 20231^T^. Moreover, using a 79% dDDH threshold as previously introduced (26), six of identified *S. aureus* (GCF_1, GCF_11, GCF_13, GCF_14, GCF_37, and GCF_38) were subclustered within the *S. aureus* population **Figure 3**. The resulting species and subspecies clusters and the taxonomic identification of the uncharacterized strains are listed in **Table S3**.

**Figure 3.**
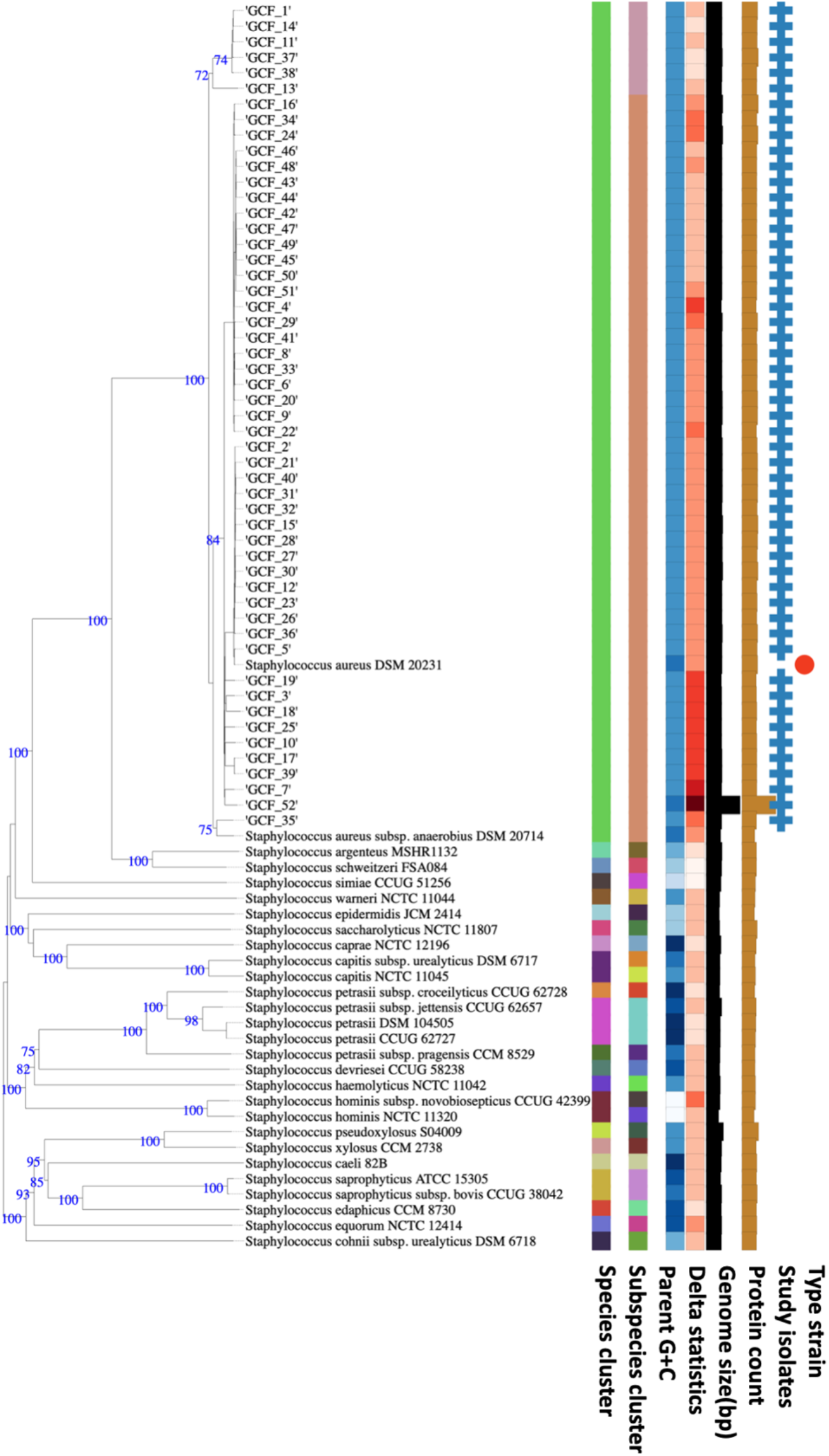
Tree inferred with FastME 2.1.6.1 from GBDP distances calculated from genome sequences. The branch lengths are scaled in terms of GBDP distance formula *d_5_*. The numbers above branches are GBDP pseudo-bootstrap support values > 60 % from 100 replications, with average branch support of 36.0 %. The tree was rooted at the midpoint.

### Multidrug resistance and virulence genes were detected among *S. aureus* isolates

We used a protein-focused approach, AMRFinderPlus, to predict horizontally acquired antimicrobial resistance (AMR) genes and resistance alleles due to chromosomal mutations (20). Distribution of AMR determinants varied significantly across genomes **Table S4**. A query gene is accepted as a true AMR gene if it reaches a threshold of 95% sequence identity and 95% coverage when compared to the database. In Staphylococcus, two mechanisms confer penicillin resistance: the production of *blaZ*-encoded beta-lactamase, which inactivates the beta-lactam ring of penicillin by hydrolyzing the peptide bonds, and the *mecA*-encoded penicillin-binding protein PBP2a which has a lower affinity for all beta-lactam antibiotics (27). We detected *blaZ* in 34 isolates representing 65.38% of the *S. aureus* population, and the *mecA* gene in 21 genomes representing 40.38% of the *S. aureus* population. Tetracyclines are broad-spectrum antibiotics used to treat many bacterial infections, and most tetracycline-resistant bacteria acquired tetracycline-resistant genes (*tet*). Two principal mechanisms of tetracycline resistance were identified in *S. aureus*: active efflux resulting from the acquisition of plasmid-located *tetK* and *tetL* genes or chromosomal-encoded Tet38 and ribosomal protection by elongation factor-like proteins that are encoded by chromosomal or transposon tetM or tetO determinants. The *tet38* gene was found in 51 isolates (98.07%), while *tetL* and *tetM* were detected in one (1.92%) and two (3.84%) genomes, respectively. Mupirocin-resistant isoleucine-tRNA ligase gene (*mupA)* was detected in one isolate (1.92%). Fosfomycin (FOM) is an antibiotic that inhibits UDP-N-acetylglucosamine enolpyruvyl transferase (MurA), an enzyme involved in the synthesis of the N-acetylmuramic acid, the essential component of peptidoglycan. Bacterial resistance to fosfomycin is due to chromosomal mutations and the expression of plasmid-encoded fosfomycin-modifying enzymes. Mutations in *murA* have been shown to lower fosfomycin affinity for MurA (28). Additionally, fosfomycin intake may be decreased in the presence of mutations in *glpT* and/or *uhpT* that encode bacterial transport systems for fosfomycin (29, 30). Finally, fosfomycin activity may be inhibited by the catalytic activity of FosA, FosB, FosC, and FosX, respectively (31–33). Among all plasmid-mediated fosfomycin resistance genes, only the *fosB* gene was identified in Staphylococcus species (34). We detected four distinct point mutations in the *murA* gene. TypeII_*murA*_ (Gly257Asp) was found in 13 isolates, TypeIII_*murA*_ (Asp278Glu) was found in 5 isolates, TypeIV_*murA*_ (Glu291Asp) was found in 7 isolates, and TypeVI_*murA*_ (Thr396Asn) was found in 2 isolates. Moreover, five genomes contained TypeIV_*glpT*_ (Val213Ile). The *FosB* gene was detected in 46 genomes. Many genomes carry genes that encode resistance against aminoglycosides (*aadD1*, *aph (3′)-IIIa*, *ant (9′)-Ia*; n = 8, 10, and 14 genomes, respectively), and streptothricin (*sat4*; n = 10 genomes). We also detected resistance determinants for macrolides (*abc-f*, *erm(A)* and *erm(C)*; n = 8, 10, and 14 genomes, respectively) Mutations in both *gyrA* (Ser84Ala; n = 1 genome and Ser84Leu; n = 13 genomes) and *parC* (Glu84Gly; n = 2 genomes, Glu84Lys; n = 1, Ser80Phe; n = 11 genomes, Ser80Tyr; n = 6 genomes) were detected which is strongly linked to reduced fluoroquinolone susceptibility (35) **Table S4**.

We identified several *S. aureus* virulence genes in the *S. aureus* population **Table S5**. Genes coding for staphylocoagulase, adenosine synthase A (*adsA*), IgG-binding protein (*Sbi*), gamma-hemolysin component A and C, alpha-hemolysin (*hla*) beta-hemolysin (*hlb*), delta-hemolysin (*hld*), capsular polysaccharide synthesis enzyme (*cap8A-G* and *cap8L-P*), iron-regulated surface determinant protein A, B, C, D, E, F and G, cell surface elastin binding protein (*ebp*), NPQTN specific sortase B, zinc metalloproteinase aureolysin (*aur*) were detected in all of the *S. aureus* isolates. Genes coding for gamma-hemolysin component A (n= 50 genomes (96.15%)), Staphylokinase precursor (*sak*; n= 44 genomes (84.61%)) and complement inhibitor SCIN (*scn*; n= 46 genomes (88.46%)) were also detected.

### SCC*mec* typing, spa typing

We examined the presence and types of SCC*mec* chromosomal cassette, which may promote the mobilization and distribution of *mec*A and other AMR genes in Staphylococcus. SCC*mec* elements are highly variable in structural organization and gene content. Still, they are classified mainly based on the *ccr* and *mec* gene complexes, the key elements of the cassette responsible for SCC*mec* integration and excision, and the beta-lactam resistance phenotype, respectively. To date, in *S. aureus* a total of 13 SCC*mec* types and various subtypes were described. We identified two known types of 21 *S. aureus* genomes carrying SCC*mec* (Type II, n=9 genomes (42.85%); Type IV, n=11 genomes (52.38%)). **Table.1**.

**Table.1:** Predicted SCC *mec* element in *S. aureus* population

| No. | Sample | Predicted SCCmec element | %template coverage |
| --- | --- | --- | --- |
| 1 | GCF_2 | SCCmec_type_IVa(2B) | 88.91% |
| 2 | GCF_4 | SCCmec_type_II(2A) | 99.75% |
| 3 | GCF_9 | SCCmec_type_II(2A) | 99.29% |
| 4 | GCF_12 | SCCmec_type_IVa(2B) | 87.45% |
| 5 | GCF_15 | SCCmec_type_IVa(2B) | 87.81% |
| 6 | GCF_16 | SCCmec_type_II(2A) | 99.09% |
| 7 | GCF_20 | SCCmec_type_II(2A) | 99.59% |
| 8 | GCF_21 | SCCmec_type_IVa(2B) | 86.78% |
| 9 | GCF_23 | SCCmec_type_IVa(2B) | 87.73% |
| 10 | GCF_24 | SCCmec_type_II(2A) | 99.60% |
| 11 | GCF_26 | SCCmec_type_IVa(2B) | 88.63% |
| 12 | GCF_27 | SCCmec_type_IVa(2B) | 88.11% |
| 13 | GCF_28 | SCCmec_type_IVa(2B) | 87.33% |
| 14 | GCF_29 | SCCmec_type_II(2A) | 99.27% |
| 15 | GCF_30 | SCCmec_type_IVa(2B) | 87.56% |
| 16 | GCF_31 | SCCmec_type_IVa(2B) | 87.17% |
| 17 | GCF_32 | SCCmec_type_IVa(2B) | 87.29% |
| 18 | GCF_33 | SCCmec_type_II(2A) | 99.18% |
| 19 | GCF_34 | SCCmec_type_II(2A) | 99.38% |
| 20 | GCF_36 | SCCmec_type_IVg(2B) | 79.41% |
| 21 | GCF_38 | SCCmec_type_II(2A) | 51.08% |

Spa typing of *S. aureus* isolates revealed fifteen different spa types (t002, t008, t189, t091, t535, t227, t017, t338, t723, t4359, t4601, t1055, t7180, t4238, and t3240) in 28 genomes (53.84%) **Table S6**. The spa types t002, t008, t189 were common among ten strains (35.71%), four strains (14.28%), and two strains (7.14%), respectively.

### A core genome single-nucleotide-polymorphism (SNP) phylogeny

Core genome SNPs-based phylogeny has a higher discriminatory speciation power compared to 16S rRNA-based phylogenetic clustering. Variants were called using snippy, which wraps BWA-MEM, SAMtools, SnpEff, and Freebayes. Core SNPs were identified as variant sites present in all samples and extracted at default settings with snippy. *S. aureus* NCTC 8325 was used as a reference genome for variant annotations. GCF-13 (ST508) was overly distinct in its high total number of SNPs, both inside and outside the boundaries of recombination blocks. GCF_45, GCF_51, GCF_49, GCF_4, GCF_20, GCF_33, GCF_2, GCF_26, GCF_27, GCF_3 and GCF_1 had no SNPs **Figure.4**.

**Figure.4:**
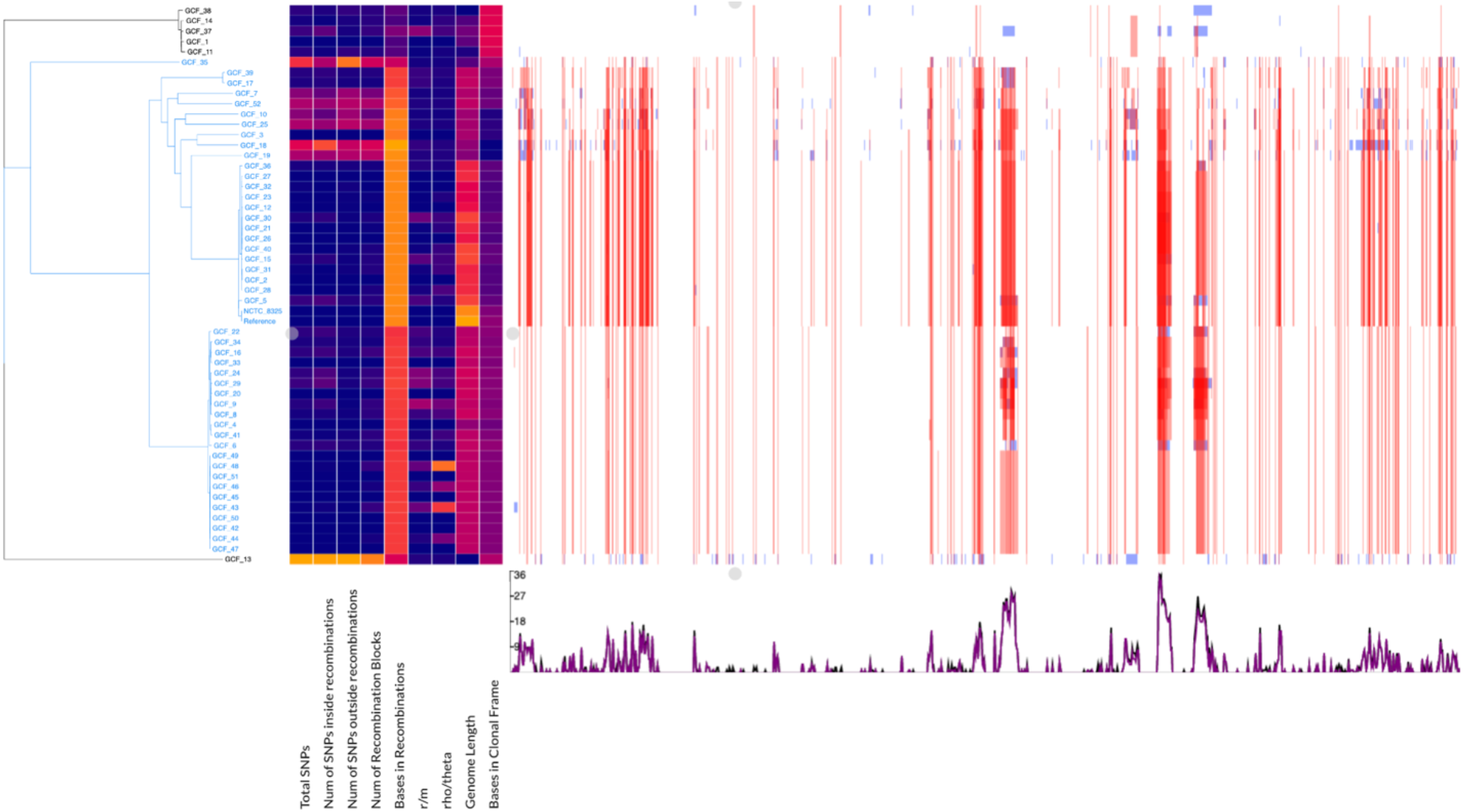
Core genome SNPs-based phylogenetic tree and genome-wide recombination hotspots.

## Discussion

A total of 54 species of staphylococcus are publicly accessible in Genbank, where the genomes of the same species are grouped under a single ID. There are, however, 253 unknown genomes (ID:13533) annotated as isolates in a staphylococcus population, which are significantly different from currently recognized species. In this study, using a genomic approach, we identified these genomes at the species level. About 182 uncharacterized Staphylococcus genomes were identified at the species level based on MLST, including 32 *S. lugdunensis* genome sequence, thus doubling the number of the publicly accessible *S. lugdunensis* genome sequence in Genbank. MLST identified five species, *S. aureus*, *S. epidermidis*, *S. lugdunensis*, *S. haemolyticus* and *S. hominis*, from uncategorized genome sequences in the NCBI database.

We identified several *S. aureus* virulence and antimicrobial resistance genes in the *S. aureus* population. There are six isolates (GCF_1, GCF_11 (ST30), GCF_13, GCF_14 (ST30), GCF_37 (ST30), and GCF_38) were subclustered close to each other within the *S. aureus* population, having similar preference in recombining loci. These isolates, except GCF_13, are of sequence type 30 and belong to clonal complex 30. However, GCF-13 (ST508) was overly distinct in its high total number of SNPs, both inside and outside the boundaries of recombination blocks.

Fifty-six isolates were not typed by MLST (20.94%). These isolates may belong to staphylococcus species that do not have MLST databases such as *Staphylococcus schleiferi* and *Staphylococcus cornubiensis,* which calls attention to the need for creating MLST databases for these species.

We used a protein-focused approach, AMRFinderPlus, to predict horizontally acquired antimicrobial resistance (AMR) genes and resistance alleles due to chromosomal mutations for several reasons. First, protein annotation and similarity comparisons to both reference proteins and the use of Hidden Markov models (HMMs) with appropriate cutoffs may help to decide whether the gene is in frame and of the correct length. In contrast, a nucleotide approach can miss nonsense mutations. Second, the AMR function is encoded by the protein sequence. Even changes in single amino acid can alter resistance phenotypes considerably, and that variation should be explicitly captured. Thirdly, discordance between nucleotides and protein sequences can lead to alleles misassignment and, thus, the possibility of the incorrect prediction of AMR (20).

In summary, our study 182 uncharacterized Staphylococcus genomes were identified, including 52 *S. aureus* genome sequences. Comparative genomics of the identified *S. aureus* isolates will give additional insights and enhances our knowledge of *S. aureus* species’ evolution.

## Notes

### Competing Interest Statement

The authors have declared no competing interest.

